# Uncovering Hidden Phenotypes in NEX-Cre Mice: Behavioral and Cellular Alterations Demand Re-Evaluation of a Widely Used Transgenic Line

**DOI:** 10.1101/2025.05.21.655260

**Authors:** Kim Renken, Olivia Andrea Masseck

## Abstract

Transgenic mouse strains are essential tools in neuroscience, enabling targeted genetic manipulations to investigate brain function and neurological diseases. The NEX-Cre mouse line, which targets glutamatergic principal neurons in the neocortex and hippocampus by expressing Cre-recombinase under the NEX (NeuroD6) promotor, has been widely used for conditional gene manipulation. Contrary to previous reports suggesting no behavioral and histological abnormalities in NEX-Cre mice, our study reveals distinct behavioral and cellular phenotypes. Behavioral analyses indicate reduced anxiety-like behavior, anhedonia, and increased locomotor activity in NEX-Cre (-/-) mice. Additionally, SVM analysis uncovered subtle strain-specific and genotype-specific behavioral traits across all NEX-Cre genotypes relative to the commonly used C57BL/6J mouse strain. Histological analyses of Golgi-Cox-stained brain slices revealed alterations in dendritic spine density across key brain regions, including the caudate putamen, hippocampal CA1, nucleus accumbens core region, lateral septum, and medial prefrontal cortex. These findings highlight significant inter-and intra-strain variability, emphasizing the importance of careful characterization of transgenic models and the need for appropriate control groups and experimental designs to ensure the reliability and validity of studies utilizing Cre-Driver lines.

## Introduction

Transgenic mouse models are indispensable tools for investigating the highly complex mechanisms underlying central nervous system (CNS) functions. They enable precise genetic manipulations to study specific molecular pathways and cellular processes *in vivo*, providing critical insights into neurodevelopment, behavior, and disease. Transcription factors are essential regulators of neurodevelopment, orchestrating processes like neuronal differentiation, mitochondrial function, and neuroprotection. The transgenic NEX-Cre mouse line expresses Cre recombinase under the control of the NEX promoter, specifically targeting glutamatergic principal neurons in the neocortex and hippocampus ^1^. This transgenic mouse line serves as a tool for behavioral research and conditional manipulation of target genes in pyramidal neurons of the dorsal telencephalon to study cortical development, learning and memory. The NEX gene was originally identified as NEX-1 ^2^ but later became known under several other names, including MATH-2, ATOH-2 or NeuroD6. Although the NEX gene is now commonly referred to as NeuroD6, we retain the term’NEX-Cre’ to align with the original nomenclature under which this transgenic mouse line was first described ^1^ and widely recognized in the scientific literature.

As a member of the NeuroD family of basic Helix-loop-Helic (bHLH) transcription factors, NEX plays a key role in neuronal differentiation, mitochondrial dynamics, and synaptic function. The NeuroD family is characterized by overlapping spatiotemporal expression patterns in the dorsal telencephalic neuroepithelium during early development and a certain degree of functional redundancy is assumed among these factors ^3–5^. The assumption of a functional redundancy between the members of the NeuroD family in combination with the specific expression of NEX in forebrain glutamatergic principal neurons and the non-lethal effects of NEX deficiency made NEX to an ideal target gene for the generation of NEX-Cre animals ^1,3^. However, the knock-in of Cre-recombinase into the NEX locus by homologous recombination in embryonic stem cells renders the NEX gene permanently non-functional, raising concerns about the potential developmental consequences. NEX regulates anti-apoptotic factors, molecular chaperones, and reactive oxygen species metabolism, which are essential for neuronal survival ^6–8^. NEX contributes to cytoskeletal remodeling, a process essential for dendritic spine formation, synaptic plasticity, and overall neuronal health ^9^. In addition to its expression in glutamatergic principal neurons of the neocortex and hippocampus ^1,3^, it is also expressed in a subpopulation of midbrain dopaminergic (mDA) neurons in the ventral tegmental area (VTA) which project to the nucleus accumbens shell ^10–13^.

Optogenetic activation of NEX-expressing mDA neurons in the VTA induces dopamine release, glutamatergic postsynaptic responses, and real-time place preference ^13^, while silencing impairs dopamine release and leads to behavioral abnormalities, such as increased consummatory behavior without changes in motivation for sucrose rewards ^14^. While these data suggest a role for NEX in behavioral modulation, the precise impact of genetic modifications in NEX-Cre animals on their behavior remains unclear. Initial reports about the NEX-Cre mice indicated no apparent histological or behavioral abnormalities ^1,3^. While some studies reported reduced anxiety-like behavior, mild anhedonic-like behavior, motor abnormalities, and learning deficits, others did not observe such abnormalities ^15–17^.

Transgenic mouse strains, including NEX-Cre mice, often exhibit behavioral or cellular abnormalities that might complicate data interpretation. These issues may arise from gene overexpression, gene knockout or off-target effects, which can unexpectedly affect protein levels, cellular functions, and downstream pathways and might often leading to pleiotropic effects and contributing to the development of neurodevelopmental and neurodegenerative diseases ^18^. Insertional mutagenesis and compensatory mechanisms may further lead to phenotypic changes and even the genetic background of the strain introduces variability into experimental outcomes. Cre-recombinase itself has been associated with behavioral changes, such as hyperactivity and impulsivity, as well as developmental defects ^19,20^.Therefore, thorough characterization of transgenic mouse lines is critical to prevent misinterpretation or incorrect attribution of observed phenotypes to specific genetic modifications. This is particularly important when inherent behavioural abnormalities cannot be excluded and could potentially interfere with experimental results.

In the present study, the behaviour and cellular architecture of all NEX-Cre genotypes, including heterozygous (wt/-) and homozygous (-/-) mutants as well as wild-type littermates (wt/wt), were characterized and compared to the commonly used C57BL/6J mouse strain. Our findings reveal clear behavioral abnormalities in NEX-Cre (-/-) mice, including reduced anxiety-like behavior, hyperlocomotion, motor deficits, and spine density abnormalities in key brain regions such as the nucleus accumbens, hippocampus, and caudate putamen. Spine density abnormalities were also observed in NEX-Cre (wt/wt) and NEX-Cre (wt/-) mice.

## Results

### NEX-Cre mice exhibit reduced anxiety-related behavior and increased locomotor activity

A comprehensive series of experiments was performed to characterize and compare the behavioral profiles of C57BL/6J, NEX-Cre (wt/wt), NEX-Cre (wt/-), and NEX-Cre (-/-) mice (**Figure 1A**). To assess anxiety-related behavior and general locomotion, three distinct behavioral paradigms were employed: the Elevated Plus Maze (EPM), Open Field Test (OFT), and Novelty-Suppressed Feeding Test (NSFT). Among these, the NSFT primarily serves as a measure of anxiety-like behavior, but it also provides insights into hunger, motivation to eat, and the animal’s ability to overcome the conflict between the drive to feed and the aversiveness of a novel environment.

**Figure 1:**
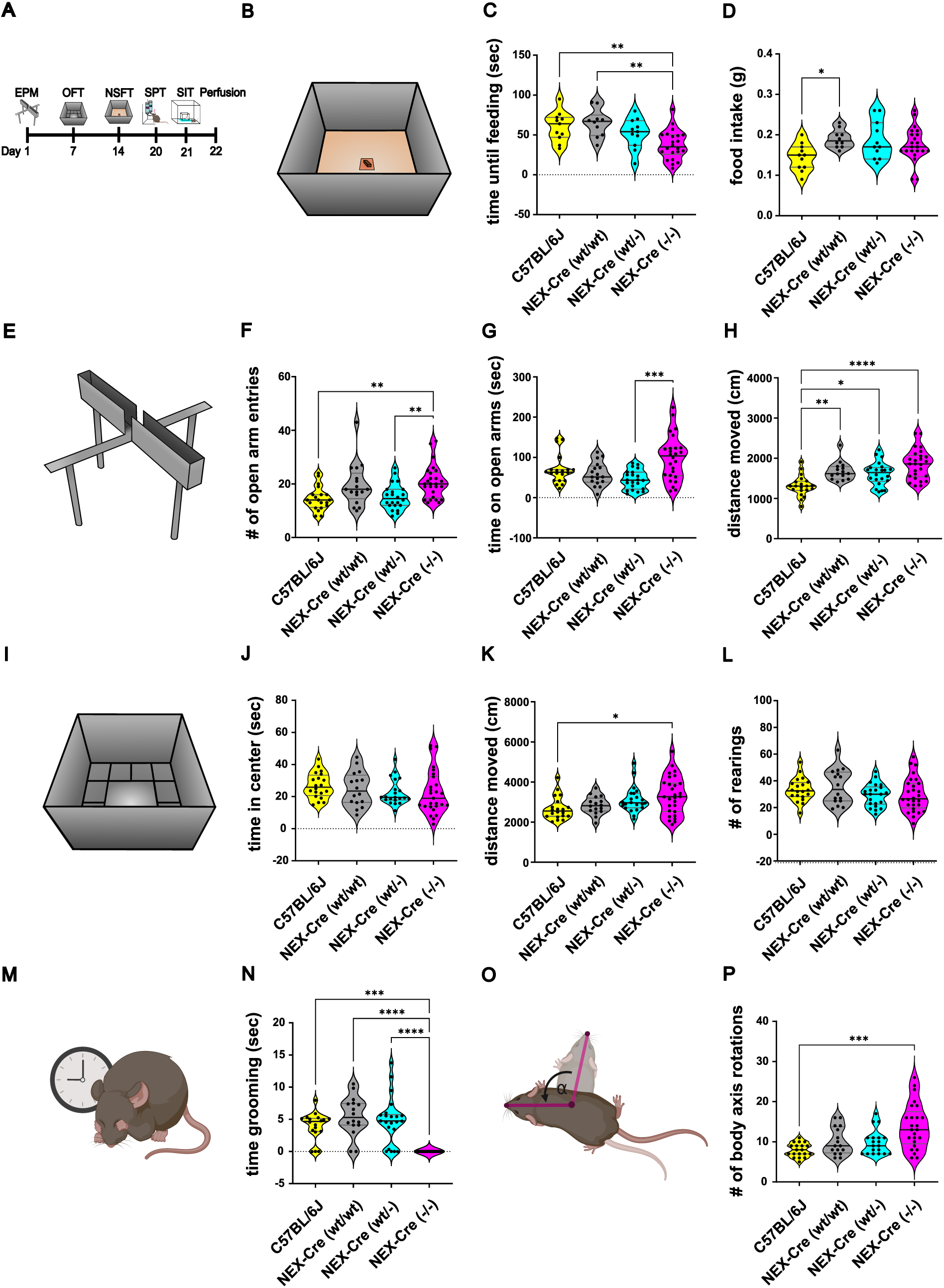
**Altered anxiety-like behavior and locomotor activity in NEX-Cre mice compared to C57BL/6J controls in the EPM, OFT, and NSFT. (**A) Timeline of behavioral experiments. (B) Schematic overview of the NSFT. (C) In NSFT, NEX-Cre (-/-) showed a significantly decreased time until feeding compared to C57BL/6J mice and NEX-Cre (wt/wt) (Ordinary one-way ANOVA: F (3,50) = 6.93, ***p = 0.0005; Tukey‘s post hoc: C57BL/6J vs NEX-Cre (-/-) **p = 0.0065, NEX-Cre (wt/wt) vs NEX-Cre (-/-) **p = 0.0014). Sample sizes: C57BL/6J: n = 11, NEX-Cre (wt/wt): n = 10, NEX-Cre (wt/-): n = 11, NEX-Cre (-/-): n = 22. (D) During the feeding phase of the NSFT, NEX-Cre (wt/wt) ate significantly more food compared to the C57BL/6J mice (Ordinary one-way ANOVA: F (3,50) = 3.128, p = 0.0338; Tukey‘s post hoc: C57BL/6J vs NEX-Cre (wt/wt) *p = 0.044). Sample sizes: C57BL/6J: n = 11, NEX-Cre (wt/wt): n = 10, NEX-Cre (wt/-): n = 11, NEX-Cre (-/-): n = 22. (E) Schematic Overview of the EPM. (F) Significantly increased open arm entries of the NEX-Cre (-/-) compared to the NEX-Cre (wt/-) and the C57BL/6J mice in the EPM (Kruskal Wallis ANOVA: H (3) = 17.02, p = 0.0007; Dunn‘s post hoc: C57BL/6J vs NEX-Cre (-/-) **p = 0.0027, NEX-Cre (wt/-) vs NEX-Cre (-/-) **p = 0.0086). Sample sizes: C57BL/6J: n = 20, NEX-Cre (wt/wt): n =1 7, NEX-Cre (wt/-): n = 22, NEX-Cre (-/-): n = 26. (G) NEX-Cre (-/-) spend significantly more time on the open arms of the EPM compared to the NEX-Cre (wt/-) mice (Kruskal Wallis ANOVA: H (3) = 16.17, p = 0.001; Dunn‘s post hoc: NEX-Cre (wt/-) vs NEX-Cre (-/-) ***p = 0.0005). Sample sizes: C57BL/6J: n = 20, NEX-Cre (wt/wt): n = 17, NEX-Cre (wt/-): n = 22, NEX-Cre (-/-): n = 26. (H) Increased locomotor activity of NEX-Cre mice in the EPM compared to C57BL/6J mice (Kruskal Wallis ANOVA: H (3) = 26.5, p < 0.0001; Dunn‘s post hoc: C57BL/6J vs NEX-Cre (wt/wt) **p =0.0046, C57BL/6J vs NEX-Cre (wt/-) *p = 0.0195, C57BL/6J vs NEX-Cre (-/-) ****p < 0.0001). (I) Schematic overview of the OFT. Sample sizes: C57BL/6J: n = 20, NEX-Cre (wt/wt): n = 15, NEX-Cre (wt/-): n = 22, NEX-Cre (-/-): n = 26. (J) All mice spend similar time in the center field in the OFT (Kruskal Wallis ANOVA: H (3) = 4.667, p=0.1979). Sample sizes: C57BL/6J: n = 20, NEX-Cre (wt/wt): n = 16, NEX-Cre (wt/-): n = 21, NEX-Cre (-/-): n = 27. (K) Increased locomotor activity of NEX-Cre (-/-) mice in the OFT compared to C57BL/6J mice (Kruskal Wallis ANOVA: H (3) = 9.572, p = 0.0226; Dunn‘s post hoc: C57BL/6J vs NEX-Cre (-/-) *p=0.0247). Sample sizes: C57BL/6J: n = 20, NEX-Cre (wt/wt): n = 16, NEX-Cre (wt/-): n = 22, NEX-Cre (-/-): n = 28. (L) All mice showed similar rearing behavior in the OFT (Ordinary one-way ANOVA: F (3,82) =1.495, p = 0.2221). Sample sizes: C57BL/6J: n = 20, NEX-Cre (wt/wt): n = 16, NEX-Cre (wt/-): n = 22, NEX-Cre (-/-): n =28. (M) Schematic representation of grooming behavior. (N) NEX-Cre (-/-) spend significantly less time grooming compared to NEX-Cre littermates and C57BL/6J mice during the OFT (Kruskal Wallis ANOVA: H (3) = 35.22, p < 0.0001; Dunn‘s post hoc: C57BL/6J vs NEX-Cre (-/-) ***p = 0.0003, NEX-Cre (wt/wt) vs NEX-Cre (-/-) ****p<0.0001, NEX-Cre (wt/-) vs NEX-Cre (-/-) ****p<0.0001). Sample sizes: C57BL/6J: n = 18, NEX-Cre (wt/wt): n = 16, NEX-Cre (wt/-): n = 22, NEX-Cre (-/-): n = 21. (O) Schematic representation of circling behaviour tracking, where body axis rotations are determined based on the accumulation of rotation angles between the axes from the center point to the nose point. This method monitors micro-rotations of animals turning around their own axis. A rotation is recorded when the cumulative angle of rotation (α) exceeds 360°. (P) During OFT, NEX-Cre (-/-) showed significantly increased circling behavior compared to C57BL/6J mice (Kruskal Wallis ANOVA: H (3) = 17.99, p = 0.0004; Dunn‘s post hoc: C57BL/6J vs NEX-Cre (-/-) ***p = 0.0002). Sample sizes: C57BL/6J: n = 19, NEX-Cre (wt/wt): n = 16, NEX-Cre (wt/-): n = 19, NEX-Cre (-/-): n = 25. The data is presented as individual data points on violin plot. The solid black line indicates the median, while the grey lines represent the 25th and 75th percentiles.

In the NSFT, NEX-Cre (-/-) mice initiated food consumption significantly faster than both NEX-Cre (wt/wt) and C57BL/6J mice (**Figure 1C**), suggesting a reduced anxiety-like phenotype. Interestingly, NEX-Cre (wt/wt) mice consumed significantly more food than C57BL/6J controls (**Figure 1D**), which may indicate an alteration in consummatory behavior or metabolic regulation in the NEX-Cre (wt/wt) line.

Additional locomotor parameters, including total distance moved and mean velocity, are shown in **Figure S1**. NEX-Cre (-/-) mice traveled significantly shorter distances compared to both NEX-Cre (wt/wt) and NEX-Cre (wt/-) mice. In contrast, NEX-Cre (wt/wt) and NEX-Cre (wt/-) animals exhibited greater locomotor activity than C57BL/6J mice. Furthermore, NEX-Cre (-/-) mice displayed significantly lower mean velocity compared to both C57BL/6J and NEX-Cre (wt/-) mice.

Similarly, in the Elevated Plus Maze (EPM), NEX-Cre (-/-) mice entered the open arms significantly more frequently than both NEX-Cre (wt/-) and C57BL/6J mice (**Figure 1F**) and spent significantly more time in the open arms compared to NEX-Cre (wt/-) mice (**Figure 1G**). These findings are consistent with the NSFT results, further supporting a phenotype characterized by reduced anxiety-like behavior and enhanced exploratory drive. Additionally, all NEX-Cre genotypes exhibited significantly greater locomotor activity than C57BL/6J controls, as indicated by the increased total distance traveled during the EPM (**Figure 1H**). This observation suggests a general trend toward hyperactivity or elevated baseline locomotion in NEX-Cre mice, irrespective of genotype.

NEX-Cre (-/-) mice exhibited particularly striking behaviors in the EPM, characterized by impulsivity and risk-taking. During exploration of the open arms, these mice frequently slipped with their hind paws—even while running in a straight line—and often displayed repetitive and erratic movement patterns, such as circling behavior on the open arms or the central platform. Their uncoordinated turning and frequent slipping occasionally resulted in partial or complete falls from the maze. Notably, these behaviors persisted across the testing session, with no apparent signs of habituation or corrective adaptation over time.

Quantitative analysis revealed that NEX-Cre (-/-) mice were significantly more likely to slip their hind paws off the open arms compared to NEX-Cre (wt/wt), NEX-Cre (wt/-), and C57BL/6J mice (**Figure S2**). While hind paw slips were occasionally observed in C57BL/6J, NEX-Cre (wt/wt), and NEX-Cre (wt/-) mice, these events typically occurred only within the first few seconds after placement on the maze and were followed by rapid acclimation and confident locomotion. Among NEX-Cre (wt/-) mice, behavioral variability was observed—some individuals showed frequent slipping, while others navigated the maze with ease.

Additional parameters, including mean velocity, locomotor trajectories, and detailed time spent in the open arms, closed arms, and central platform, are presented in **Figure S2**. Collectively, these findings further emphasize the phenotype of reduced anxiety-like behavior, hyperactivity, and impaired motor coordination in NEX-Cre (-/-) mice.

In the OFT, all groups spent a similar amount of time in the center field (**Figure 1J**), and exhibited comparable frequencies of rearing behavior (**Figure 1L**), indicating no differences in anxiety-like or exploratory behavior between the groups. However, increased locomotor activity was observed in NEX-Cre (-/-) mice compared to C57BL/6J mice, with NEX-Cre (wt/wt) and NEX-Cre (wt/-) mice showing a slight trend toward increased locomotor activity (**Figure 1K**). Notably, NEX-Cre (-/-) mice exhibited striking behavioral abnormalities during the OFT. They did not display any self-grooming behavior during the 5-minute test period, in contrast to the other groups (**Figure 1N**). This suggests a potential correlation between either the gene dosage of Cre recombinase or the NEX knockout and self-grooming behaviour. Additionally, similar to the EPM, NEX-Cre (-/-) mice demonstrated prominent circling behavior, rotating around their own axis significantly more often than C57BL/6J mice (**Figure 1P**). Additional details on behavioral parameters in the OFT are presented in **Figure S3**, including measurements of velocity, zone entries (border, corner, and center), corresponding latencies, as well as ethologically relevant behaviors such as rearing, circling, and grooming. NEX-Cre (-/-) mice entered the border zone significantly more frequently than both NEX-Cre (wt/wt) and C57BL/6J mice and spent more time in the border zones compared to C57BL/6J animals

(**Figure S3C, G**). Moreover, both NEX-Cre (-/-) and NEX-Cre (wt/-) mice spent less time rearing than C57BL/6J controls (**Figure S3L**), suggesting a reduction in vertical exploration.

Collectively, these findings support the notion of altered motor activity in NEX-Cre (-/-) mice and highlight changes in sequential movement patterns and the presence of stereotyped behaviors. These alterations further distinguish the NEX-Cre (-/-) behavioral phenotype from that of wild-type and heterozygous counterparts.

In summary, results indicate reduced anxiety-related behavior and increased locomotor activity in the NEX-Cre groups compared to the C57BL/6J controls. The gradual progression of the measured behaviors suggests that the genetic background of the NEX animals influences their behavior. However, in contrast to the NSFT and EPM, the OFT did not reveal any clear indications of altered anxiety-like behavior in NEX-Cre (-/-) mice. This discrepancy may reflect the differing sensitivities and contextual demands of the respective behavioral paradigms. Together, these findings underscore the complex interplay between genetic alterations, locomotor activity, and anxiety-related behaviors, and highlight the importance of employing multiple complementary assays to capture the multifaceted nature of behavioral phenotypes.

### NEX-Cre mice show normal social and mild anhedonic-like behavior

To assess social behavior in NEX-Cre (wt/wt), NEX-Cre (wt/-), NEX-Cre (-/-), and C57BL/6J mice, we performed the social interaction (SI) test and calculated SI ratios (**Figure 2A–C**). No significant differences in SI ratios were observed between groups (**Figure 2B**). Notably, all C57BL/6J mice exhibited SI ratios above 1, whereas some mice in the NEX-Cre groups fell below this threshold. Socialization time, defined as the percentage of time spent in the interaction zone during the interaction phase ^21^, also did not differ between groups (**Figure 2C**).

**Figure 2.**
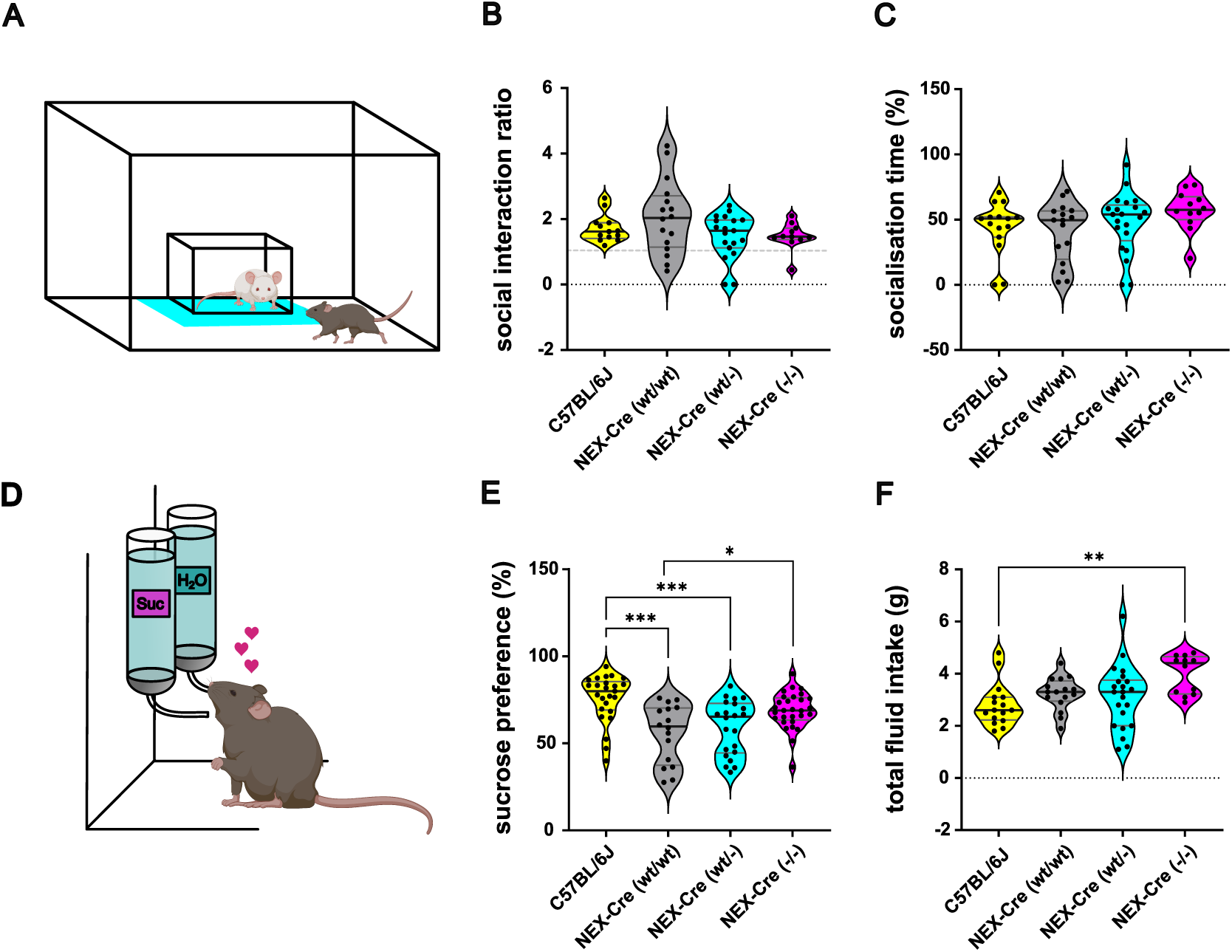
Decreased sucrose preference and normal social behavior in NEX-Cre mice. (A) Schematic overview of the social interaction test. (B) SI-ratios do not differ between the groups (Ordinary one-way ANOVA: F (3,55) = 2.061, p=0.116). Sample sizes: C57BL/6J: n = 13, NEX-Cre (wt/wt): n = 16, NEX-Cre (wt/-): n = 19, NEX-Cre (-/-): n = 11. (C) No difference between the groups in the socialization time during social interaction test (Kruskal Wallis ANOVA: H (3) = 4.941, p=0.176). Sample sizes: C57BL/6J: n = 15, NEX-Cre (wt/wt): n = 16, NEX-Cre (wt/-): n = 21, NEX-Cre (-/-): n = 12. (D) Schematic overview of the sucrose preference test. (E) Decreased sucrose preference in NEX-Cre (wt/wt) and NEX-Cre (wt/-) compared to C57Bl76J mice (Ordinary one-way ANOVA: F (3,88) = 9.376 ****p < 0.0001; Tukey‘s post hoc: C57BL/6J vs NEX-Cre (wt/wt) ****p < 0.0001, C57BL/6J vs NEX-Cre (wt/-) ***p = 0.0005, NEX-Cre (wt/wt) vs NEX-Cre (-/-) *p = 0.0174). Sample sizes: C57BL/6J: n = 25, NEX-Cre (wt/wt): n = 16, NEX-Cre (wt/-): n = 22, NEX-Cre (-/-): n = 29. (F) NEX-Cre (-/-) showed increased liquid intake compared to C57Bl76J mice (Kruskal Wallis ANOVA: H (3) = 11.99, p = 0.0074; Dunn‘s post hoc: C57BL/6J vs NEX-Cre (-/-) **p=0.0037). Sample sizes: C57BL/6J: n = 16, NEX-Cre (wt/wt): n = 16, NEX-Cre (wt/-): n = 22, NEX-Cre (-/-): n = 12.

In the sucrose preference test (SPT), NEX-Cre (wt/wt) and NEX-Cre (wt/-) mice exhibited a significantly reduced sucrose preference compared to C57BL/6J mice (median values:

C57BL/6J: 80%, NEX-Cre (wt/wt): 59.75%, NEX-Cre (wt/-): 65.33%; p = 0.0005, p < 0.0001; ***, ****), suggesting innate anhedonic-like behavior or altered reward sensitivity (**Figure 2E**). Interestingly, NEX-Cre (-/-) mice demonstrated a significantly increased sucrose preference compared to NEX-Cre (wt/wt) mice (median values: NEX-Cre (wt/wt): 59.75%, NEX-Cre (-/-): 68.89%; p = 0.0174; *), indicating heightened consummatory behavior relative to their wildtype littermates. Furthermore, during the 6-hour SPT, NEX-Cre (-/-) mice consumed significantly more sucrose and water than C57BL/6J mice (**Figure 2F**). Although the median fluid intake of NEX-Cre (wt/wt) and NEX-Cre (wt/-) mice was slightly elevated, these differences did not reach statistical significance. The increased fluid intake observed in NEX-Cre (-/-) mice may reflect an elevated metabolic rate.

The social interaction test revealed no significant group differences, although some NEX-Cre mice showed reduced social interest. NEX-Cre (wt/wt) and NEX-Cre (wt/-) mice displayed lower sucrose preference, suggesting innate anhedonia or altered reward sensitivity. In contrast, NEX-Cre (-/-) mice showed increased sucrose preference and fluid intake, indicating heightened consumption and possibly elevated metabolism.

### Classification analysis of EPM and OFT behavioral data reveal subtle but distinct behavioral differences between NEX-Cre genotypes and C57BL/6J mice

Although significant behavioral deficits were primarily observed in NEX-Cre (-/-) mice, we hypothesized that additional subtle differences might exist across genotypes that were not fully captured by traditional behavioral measures. To investigate this, we applied advanced machine learning techniques, including principal component analysis (PCA), t-distributed stochastic neighbor embedding (t-SNE), and support vector machine (SVM) classification, to uncover potential hidden behavioral patterns. Data from the OFT and EPM (**Figures 1, S2, and S3**) served as the basis for these analyses, providing a comprehensive behavioral dataset for genotype classification.

To assess whether spontaneous behavior alone is sufficient to differentiate between genotypes, we first applied PCA and t-SNE to the behavioral feature set. Neither method revealed any clear clustering or separation by genotype (**Figure 3A-B**), suggesting that the behavioral differences are subtle and not linearly separable in the high-dimensional space. First, PCA was performed to provide an initial understanding and visualization of the data structure, capturing 38% of the total variance (**Figure 3A**). While PC1 and PC2 accounted for a substantial portion of the variance, the remaining 62% indicated additional complexity requiring further exploration.To investigate local structure that might not be captured by the linear nature of PCA, t-SNE was applied as a non-linear dimensionality reduction technique

**Figure 3.**
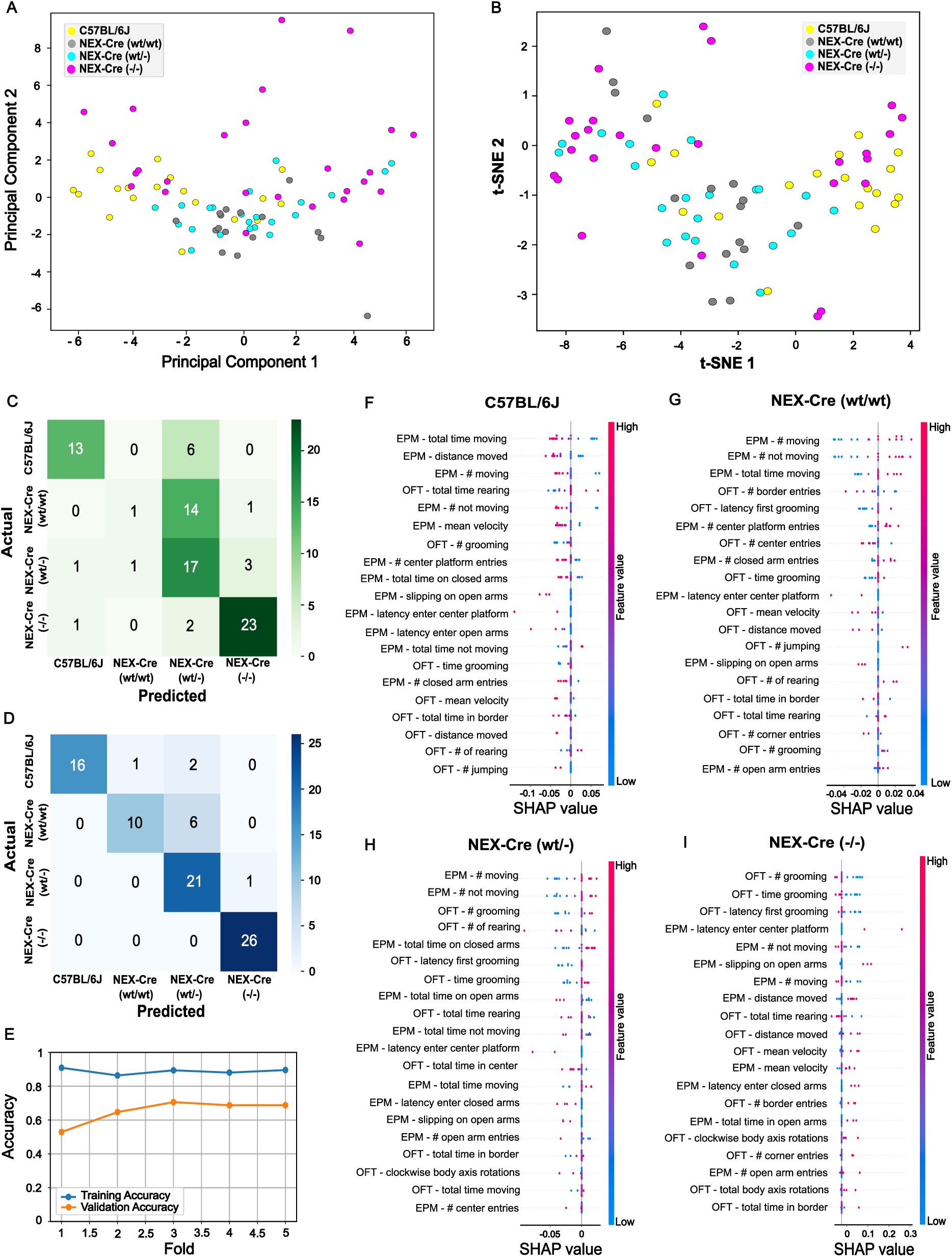
Dimensionality Reduction and Classification Analysis of Behavioral Data from EPM and OFT: PCA, t-SNE, SVM Confusion Matrix, and SHAP Value Insights. (A) Visualization of the dataset after transformation and dimensionality reduction using PCA, highlighting the primary patterns and capturing 38% of the variance in the dataset. (B) Visualization of the dataset preprocessed by PCA and then transformed and reduced to two dimensions using t-SNE, with a silhouette score of-0.092 indicating strong cluster overlap, and a trustworthiness score of 0.85 reflecting a high level of preservation of local data structure. The silhouette score of - 0.065 suggest strong overlap of the groups (C) Aggregated confusion matrix using 5-fold stratified cross-validation reached a mean accuracy of ∼65% and a F1-score of ∼ 0.6. (D) Confusion matrix of the entire dataset, mean accuracy 88%. (E) Model training and validation accuracy. (D-G) Beeswarm plot visualization of SHAP values derived from the SVM model, illustrating the contribution of features to the prediction of the groups: C57BL/6J (F), NEX-Cre (wt/wt) (G), NEX-Cre (wt/-) (H), and NEX-Cre (-/-) mice (I). Each row represents a feature, ranked from top to bottom by the mean absolute SHAP value. Individual dots in each row represent SHAP values for that feature, with color indicating the feature value according to the color bar. The position along the x-axis reflects the SHAP value, representing the feature’s impact on the model’s prediction. Sample sizes: C57BL/6J: n = 19, NEX-Cre (wt/wt): n = 16, NEX-Cre (wt/-): n = 22, NEX-Cre (-/-): n =26.

(**Figure 3B**). The method yielded a trustworthiness score of 0.85, indicating good preservation of local relationships in the high-dimensional behavioral space. However, the silhouette score of –0.065 suggested strong overlap between samples from different genotypes, indicating that no meaningful cluster separation emerged in the t-SNE space. Together, these unsupervised approaches suggest that genotype-specific behavioral patterns are not trivially or linearly separable based on the observed feature space.

Despite the lack of unsupervised structure, a SVM classifier achieved robust performance in predicting genotype from behavioral features. Using 5-fold stratified cross-validation, the model reached a mean accuracy of ∼65% and a mean F1-score of ∼0.6. The aggregated confusion matrix shows clear diagonal dominance (**Figure 3C)**, indicating consistent classification across genotypes. The model significantly outperformed chance, which was defined as the accuracy of always predicting the most frequent class (∼31.3%, Binomial test, *p* < 0.001). A confusion matrix computed on the entire dataset confirms high classification accuracy (mean accuracy 0.88) (**Figure 3D**). Model training and validation accuracy were highly consistent across folds, indicating that the classifier generalizes well and is not overfitting **(Figure 3E**).

We next asked which behavioral features drive classification for each genotype. Using SHAP (SHapley Additive exPlanations), we visualized class-wise feature importances based on the final fold of the SVM model. Each genotype exhibited distinct SHAP profiles, with different top-ranked features contributing most to prediction (**Figure 3F-I**). SHAP feature attribution revealed both shared and class-specific behavioral drivers of genotype classification (**Figure 3F-I**). For instance, the-frequency of not moving in the EPM was consistently ranked among the top features across multiple genotypes, but showed the strongest and most directional contribution to predictions of the NEX-Cre (wt/-) group. In contrast, latency to first enter the center platform during EPM was an important class-specific feature for identifying the NEX-Cre (-/-) group, but contributed negligibly to predictions in the NEX-Cre (wt/wt) group. These results suggest that the classifier draws on both shared and genotype-distinct behavioral signatures. Importantly, grooming behavior surfaced as a distinctive marker: grooming frequency was strongly weighted in the classification of NEX-Cre (-/-), consistent with the striking absence of this behaviour in this group. Similarly, circling frequency, a marker often associated with stereotypy or motor abnormalities, as well as slipping in the EPM contributed more to predictions in NEX-Cre (-/-) than in other genotypes—further supporting the model’s sensitivity to subtle motor behavioral changes.

In summary, the SVM classifier was able to reliably predict genotype from behavior with an accuracy well above chance, despite the absence of clear separability in unsupervised dimensionality reduction. SHAP analysis further uncovered distinct behavioral features that contributed to genotype classification, offering interpretable insights into subtle yet meaningful behavioral differences. These findings highlight the complexity of genotype-specific behavioral signatures and underscore the role of both strain and genetic background in shaping behavior.

### Structural alterations of dendritic spine density

Given the abnormal behavioral traits observed in NEX-Cre mice, such as anhedonic-like behavior, reduced anxiety, and increased locomotion, and considering the critical role of NEX in neuronal development and survival, we investigated dendritic spine density in key brain regions associated with emotion, reward processing, decision-making, learning, memory, motor planning and social behavior. These regions included the medial prefrontal cortex (mPFC), nucleus accumbens core region (NaC), caudate putamen (CPu), dorsal (LSD) and intermediate (LSI) lateral septum, hippocampal CA1 region (Hip CA1), and basolateral amygdala (BLA). To assess dendritic spine density, Golgi-Cox staining was performed and analyzed (**Figure 4**).

**Figure 4.**
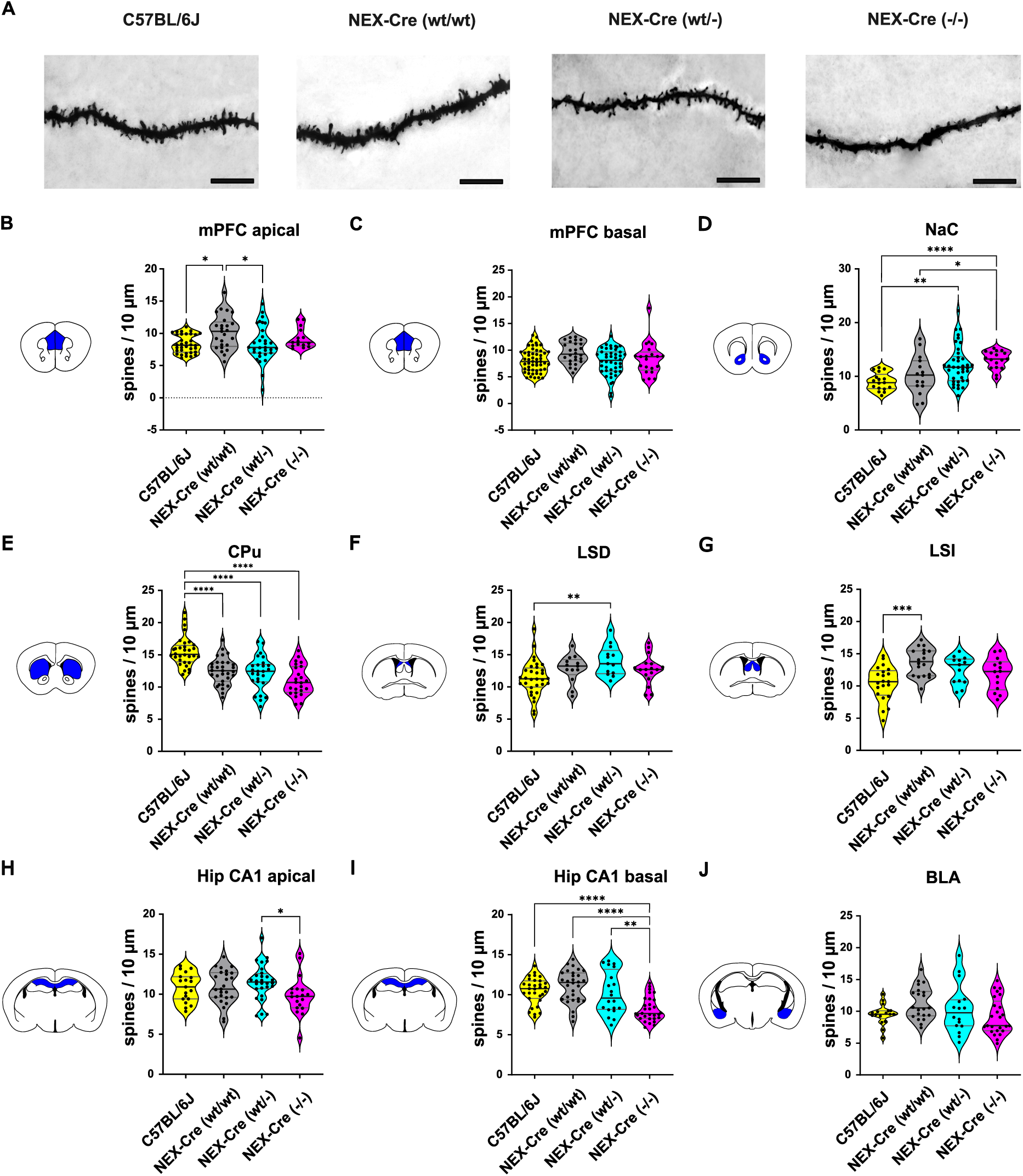
Dendritic Spine Density of NEX-Cre and C57BL/6J mice. (A) Representative images of Golgi-Cox-stained basal dendrites of pyramidal neurons in the CA1 region of the hippocampus. Scalebar 10µm. (B) Increased dendritic spine density on apical dendrites of pyramidal neurons of layer V in the PFC of NEX-Cre (wt/wt) mice compared to C57BL/6J and NEX-Cre (wt/-) mice (Kruskal Wallis ANOVA: H (3) = 10.41, p = 0.0154; Dunn‘s post hoc: C57BL/6J vs NEX-Cre (wt/wt) *p=0.0344, NEX-Cre (wt/wt) vs NEX-Cre (wt/-) *p=0.0498). Sample sizes of analyzed dendrites: C57BL/6J: n = 34, NEX-Cre (wt/wt): n = 25, NEX-Cre (wt/-): n = 31, NEX-Cre (-/-): n = 15. (C) Spine density on basal dendrites in PFC (Kruskal Wallis ANOVA: H (3) = 8.219, p= 0.0417, Dunn‘s post hoc: p-values > 0.05). Sample sizes of analyzed dendrites: C57BL/6J: n = 52, NEX-Cre (wt/wt): n = 26, NEX-Cre (wt/-): n = 44, NEX-Cre (-/-): n = 21. (D) Increased spine density on medium spiny neurons of the NaC of NEX-Cre (wt/-) and NEX-Cre (-/-) mice compared and C57BL/6J mice but also in NEX-Cre (-/-) compared to NEX-Cre (wt/wt) mice. (Kruskal Wallis ANOVA: H (3) = 22.61, p < 0.0001, Dunn‘s post hoc: C57BL/6J vs NEX-Cre (wt/-) **p=0.0029, C57BL/6J vs NEX-Cre (-/-) ****p< 0.0001, NEX-Cre (wt/wt) vs NEX-Cre (-/-) *p=0.0258). Sample sizes of analyzed dendrites: C57BL/6J: n = 17, NEX-Cre (wt/wt): n = 15, NEX-Cre (wt/-): n = 38, NEX-Cre (-/-): n = 17. (E) Decreased spine density on medium spiny neurons of the CPu in all Nex-Cre groups compared to C57BL/6J mice (Ordinary one-way ANOVA: F (3,109) = 20.46, ****p < 0.0001; Tukey‘s post hoc: C57BL/6J vs NEX-Cre (wt/wt) ****p < 0.0001, C57BL/6J vs NEX-Cre (wt/-) ****p < 0.0001, C57BL/6J vs NEX-Cre (-/-) ****p < 0.0001). Sample sizes of analyzed dendrites: C57BL/6J: n = 32, NEX-Cre (wt/wt): n = 31, NEX-Cre (wt/-): n = 26, NEX-Cre (-/-): n = 24. (F) Increased spine density on neurons the LSD in NEX-Cre (wt/-) mice compared to C57BL/6J mice (Ordinary one-way ANOVA: F (3,71) = 4.056, p = 0.0102; Tukey‘s post hoc: C57BL/6J vs NEX-Cre (wt/-) **p = 0.0074). Sample sizes of analyzed dendrites: C57BL/6J: n = 32, NEX-Cre (wt/wt): n = 13, NEX-Cre (wt/-): n = 13, NEX-Cre (-/-): n = 17. (G) Increased spine density on neurons of the LSI in NEX-Cre (wt/wt) mice compared to C57BL/6J mice (Ordinary one-way ANOVA: F (3,62) = 6.124, p = 0.001; Tukey‘s post hoc: C57BL/6J vs NEX-Cre (wt/wt) ***p = 0.0005). Sample sizes of analyzed dendrites: C57BL/6J: n = 21, NEX-Cre (wt/wt): n = 18, NEX-Cre (wt/-): n = 13, NEX-Cre (-/-): n = 14. (H) Decreased spine density on apical dendrites of pyramidal neurons in the hippocampal CA1 region in NEX-Cre (-/-) compared to NEX-Cre (wt/-) mice (Ordinary one-way ANOVA: F (3,87) = 2.43, p= 0.0705; Tukey‘s post hoc: NEX-Cre (wt/-) vs NEX-Cre (-/-) *p = 0.0412). Sample sizes of analyzed dendrites: C57BL/6J: n = 19, NEX-Cre (wt/wt): n = 25, NEX-Cre (wt/-): n = 28, NEX-Cre (-/-): n = 19. (I) Decreased spine density on basal dendrites of pyramidal neurons in the hippocampal CA1 region in NEX-Cre (-/-) compared to NEX-Cre (wt/-), NEX-Cre (wt/wt) and C57BL/6J mice (Ordinary one-way ANOVA: F (3,107) = 12.98, p<0.0001; Tukey‘s post hoc: C57BL/6J vs NEX-Cre (-/-) ****p<0.0001, NEX-Cre (wt/wt) vs NEX-Cre (-/-) ****p<0.0001, NEX-Cre (wt/-) vs NEX-Cre (-/-) **p = 0.0019). Sample sizes of analyzed dendrites: C57BL/6J: n = 31, NEX-Cre (wt/wt): n = 30, NEX-Cre (wt/-): n = 20, NEX-Cre (-/-): n = 30. (J) No differences in the dendritic spine density on neurons of the BLA between the groups (Ordinary one-way ANOVA: F (3,75) = 2.427, p=0.0712). Sample sizes of analyzed dendrites: C57BL/6J: n = 17, NEX-Cre (wt/wt): n = 21, NEX-Cre (wt/-): n = 16, NEX-Cre (-/-): n = 25.

In the mPFC, NEX-Cre (wt/wt) mice showed increased spine density on apical dendrites compared to C57BL/6J and NEX-Cre (wt/-) mice, with no differences observed on basal dendrites (**Figures 4B, 4C**). This unexpected increase suggests that external factors or subtle genetic variations may affect neural architecture in NEX-Cre (wt/wt) mice. In the NaC, spine density was elevated in NEX-Cre (-/-) mice compared to C57BL/6J and NEX-Cre (wt/wt) mice, and in NEX-Cre (wt/-) mice compared to C57BL/6J mice (**Figure 4D**), indicating possible gene dosage effects of Cre recombinase or NEX deletion. In the CPu, all NEX-Cre groups exhibited reduced spine density relative to C57BL/6J mice (**Figure 4E**). In the LSD and LSI, spine density was increased in NEX-Cre (wt/-) and NEX-Cre (wt/wt) mice, respectively, compared to C57BL/6J mice (**Figures 4F, G**).In the hippocampal CA1 region, NEX-Cre (-/-) mice displayed reduced spine density on both apical and basal dendrites, with the latter reduction more pronounced across groups (**Figures 4H, I**). In the BLA, no significant group differences were observed, although NEX-Cre (-/-) mice showed a trend toward lower spine density (**Figure 4J**).

In summary, dendritic spine density alterations were observed across specific brain regions in NEX-Cre mice. NEX-Cre (wt/wt) mice showed increased spine density in the mPFC and LSI, while all NEX-Cre groups exhibited gradual increases in the NaC, suggesting synaptic changes linked to the NEX-Cre genetic background. Conversely, all NEX-Cre groups showed reduced spine density in the CPu compared to C57BL/6J mice. NEX-Cre (-/-) mice displayed pronounced reductions in both apical and basal dendrites of the hippocampal CA1 region. No significant differences were found in the BLA. These results point to genotype-and strain-specific alterations in synaptic connectivity that may contribute to the behavioral phenotypes observed in NEX-Cre mice.

## Discussion

### Hyperlocomotion, reduced anxiety-like behavior, motor dysfunctions and anhedonic like behavior in NEX-Cre mice

NEX-Cre (-/-) mice exhibited behavioral abnormalities, including increased exploration, hyperactivity, circling behavior, and reduced self-grooming. These findings align with previous reports linking NEX knockout models to hyperlocomotion ^15,17^. In contrast, NEX-Cre (wt/-) mice showed no such abnormalities, suggesting compensation for NEX haploinsufficiency. Circling behavior, a stereotypy associated with basal ganglia dysfunction and dopaminergic dysregulation, and reduced self-grooming, a behavior modulated by forebrain circuits, indicate disrupted neural systems in NEX-Cre (-/-) mice. Given that grooming is also an important measure in models of OCD and autism spectrum disorders ^22^, its absence may reflect broader neuropsychiatric-like alterations, though motivational or competitive factors cannot be ruled out ^23^.

Reduced anxiety-like behavior in NEX-Cre (−/−) mice is consistent with previous findings using the same model ^17^, but contrasts with studies on NEX knockout mice lacking Cre expression ^15^, suggesting that Cre recombinase expression, in addition to NEX deletion, may influence the phenotype.

No major abnormalities in social interaction were observed across groups, though caution is warranted due to methodological limitations of the SI test, which may miss subtle social deficits. Future studies could employ more refined approaches, such as three-chamber tests or machine learning-based tracking (e.g., SLEAP; ^24^), to capture detailed social behavior.

In the sucrose preference test, NEX-Cre (wt/wt) and NEX-Cre (wt/-) mice showed reduced sucrose preference compared to C57BL/6J controls, suggesting an innate anhedonic-like state, a core symptom of depression characterized by diminished pleasure from rewarding stimuli. Unexpectedly, NEX-Cre (-/-) mice exhibited increased sucrose preference compared to their NEX-Cre (wt/wt) littermates, despite showing no differences relative to C57BL/6J controls. These findings align with recent studies ^14^ linking NEX-expressing midbrain dopamine neurons to consummatory behavior, suggesting that NEX-positive VTA mDA neurons play a key role in consummatory aspects of reward-related behavior ^13,14^. Consistently, NEX-Cre (-/-) mice showed increased overall fluid intake, supporting enhanced consummatory drive.

### Behavioral Classification by SVM

Despite mild and heterogeneous behavioral abnormalities in NEX-Cre mice, SVM-based classification successfully predicted genotypes using EPM and OFT data. While PCA and t-SNE did not reveal clear behavioral cluster, SVM achieved high accuracy, particularly for NEX-Cre (-/-) mice, consistent with their pronounced phenotype. Notably, the SVM also differentiated NEX-Cre (wt/wt) and (wt/-) mice from C57BL/6J controls, revealing subtle behavioral traits in these genotypes. This challenges the assumption that NEX-Cre (wt/-) mice fully compensate for haploinsufficiency.

Furthermore, distinguishing NEX-Cre (wt/wt) from C57BL/6J mice—despite identical genetic backgrounds—suggests possible maternal effects from NEX-Cre (wt/-) dams, warranting future cross-fostering studies. In contrast, genotype classification based on social interaction data failed (**Figure S4**), likely due to the limited behavioral parameters assessed.

Overall, our findings highlight motor and cognitive components in the hyperactivity-like behavior of NEX-Cre (-/-) mice and reveal genotype-specific traits in NEX-Cre (wt/wt) and (wt/-) mice, emphasizing the influence of genetic background on behavior.

### Spine density alterations in NEX-Cre mice

Given the behavioral abnormalities observed in NEX-Cre mice, we analyzed dendritic spine densities and identified structural changes in key brain regions. In the NaC, a region central to reward processing, motivation, and sucrose preference behavior, all NEX-Cre genotypes exhibited increased spine density, with the most pronounced changes in NEX-Cre (-/-) and NEX-Cre (wt/-) mice, and a moderate increase in NEX-Cre (wt/wt) mice. Increased spine density in NEX-Cre (wt/wt) and (wt/-) mice correlated with reduced sucrose preference, supporting an anhedonic-like phenotype. In contrast, NEX-Cre (-/-) mice, despite having the highest spine density, showed increased sucrose preference compared to NEX-Cre (wt/wt) littermates and no difference relative to C57BL/6J controls, suggesting that structural changes in the NaC may differentially influence reward-related behaviors across genotypes.

In the CPu, all NEX-Cre genotypes exhibited reduced spine density compared to C57BL/6J controls. As a key region involved in motor control, reward processing, habit formation, and cognitive functions, structural changes in the CPu may contribute to the circling behavior, motor deficits, and lack of grooming observed in NEX-Cre (-/-) mice ^25–27^. Interestingly, these behavioral abnormalities were absent in NEX-Cre (wt/wt) and (wt/-) mice, suggesting the presence of compensatory mechanisms that mitigate the effects of reduced spine density.

The hippocampal CA1 region showed pronounced changes, with NEX-Cre (-/-) mice exhibiting reduced spine density on both apical and basal dendrites. These structural alterations may impair synaptic plasticity, contributing to the observed hyperactivity-like behaviors and the learning and memory deficits previously reported in NEX knockout models ^15^. In wild-type animals, NEX mRNA is strongly expressed in the hippocampus, particularly in the CA1 and CA3 regions ^3^, highlighting its role in neuronal differentiation and synaptic plasticity. The observed structural impairments in NEX-Cre (-/-) mice likely reflect the consequences of NEX loss.

In the lateral septum (LS), spine density was increased in the LSI of NEX-Cre (wt/wt) mice and the LSD of NEX-Cre (wt/-) mice, whereas NEX-Cre (-/-) mice showed no significant changes. This was unexpected, given the known role of NEX in the survival of mDA neurons projecting to the LS, and may reflect compensatory stabilization of synaptic structures in the absence of NEX.

Finally, in the BLA, a key region involved in processing memory, motivation, and emotions such as fear and anger, no significant differences in spine density were detected across groups. The absence of structural changes in the BLA is consistent with the reduced anxiety-like behavior observed in NEX-Cre (-/-) mice, reflecting the complex, context-dependent and often non-linear relationship between dendritic spine alterations and behavioral outcomes.

In summary, these findings reveal region-specific synaptic alterations in NEX-Cre mice, highlighting the complexity of structural plasticity across brain regions and its potential link to behavioral phenotypes. However, whether these spine changes are causal, consequential, or compensatory remains unclear, underscoring the need for future studies investigating spine morphology, synaptic connectivity, and functional properties.

### Genetical Background and Behavior

NEX is primarily expressed in glutamatergic principal neurons, and its haploinsufficiency or knockout may impair glutamatergic pathways, contributing to hyperactivity-like behaviors. NEX is also expressed early during development (E14.5) in a subset of VTA dopaminergic (mDA) neurons projecting to the lateral septum (LS) and nucleus accumbens shell, regions involved in motor control and reward processing ^10,11,13,14,28,29^. NEX knockout studies demonstrate a reduction in VTA mDA neuron numbers, underscoring its essential role in dopaminergic development and survival ^10^.

The NEX-Cre mouse model has been widely used to investigate neuronal development, signaling, and behavior ^13,14,16,30–35^ and has facilitated the creation of advanced genetic tools, such as the DAT-P2A-Flpo line for targeting NEX-positive dopaminergic subpopulations ^11^. However, the utility of NEX-Cre mice depends on thorough characterization of their baseline phenotypes, as unrecognized behavioral or structural alterations could confound interpretations.

Importantly, knock-in of Cre recombinase itself can induce ectopic gene expression, DNA damage, and behavioral abnormalities such as hyperactivity and impulsivity, independent of the targeted gene deletion ^19,20,36–38^. These side effects are often dose-dependent and highlight the necessity of including appropriate Cre-only controls with matched allele numbers in both behavioral and cellular studies. Relying solely on behavioral normality to exclude cellular or structural abnormalities is problematic, as subtle phenotypes may be overlooked.

Although inducible Cre systems (e.g., tamoxifen-inducible) or viral strategies (e.g., CamKII promoters) offer alternatives to minimize developmental disturbances, they also have limitations. Thus, rigorous control strategies and advanced analytical approaches are crucial to accurately separate Cre-related effects from true gene-specific phenotypes, improving the reliability and interpretability of studies using Cre/loxP models.

Overall, these considerations emphasize that careful experimental design, including comprehensive phenotypic validation, is essential when using NEX-Cre mice or related models to ensure accurate conclusions about gene function and neuronal circuitry.

## Acknowledgements

We would like to thank Juliana Groß, Celina Schreiber, Frederik Piel, Hannah Urbschat and for expert technical assistance and Professor Klaus-Armin Nave and Sandra Göbbels (Max Plank Institute of Experimental Medicine, Göttingen, Germany) for the NEX-Cre mice.

## Author contribution

K.R.: conceptualization, experimental design, data acquisition and analysis, figure preparation, manuscript writing. O.M.: conceptualization,data analysis, scientific supervision, manuscript editing.

## Declaration of generative AI and AI-assisted technologies

During the preparation of this work, the author(s) used ChatGPT in order to improve language and to assist in designing the data analysis for the SVM. After using this tool or service, the author(s) reviewed and edited the content as needed and take(s) full responsibility for the content of the publication.

## Methods

### KEY RESOURCES TABLE

| REAGENT or RESOURCE | SOURCE | IDENTIFIER |
| --- | --- | --- |
| Critical commercial assays |  |  |
| FD Rapid GolgiStain™ Kit | FD NeuroTechnologies, Ellicott City | Cat# PK401 |
| Experimental models: Organisms/strains |  |  |
| C57BL/6J | The Jackson Laboratory | JAX:000664 |
| NEX-Cre<br>(or NeuroD6-Cre) | Laboratory of Prof. Nave <sup>1</sup> | N/A |
| CD-1 IGS | Charles River | CRL:022 |
| Software and algorithms |  |  |
| Python (version 3.10.1) | Python Software Foundation | https://www.python.org/ |
| EthoVision XT | Noldus (Virginia, USA) | Version: 15.0.141 |
| GraphPad Prism | GraphPad Software | Version: 9.3.1. |

## RESOURCE AVAILABILITY

### Lead contact

Further information and requests for resources and reagents should be directed to and will be fulfilled by the lead contact, Olivia Masseck

### Materials availability

This study did not generate new unique reagents.

### Data and code availability

Data reported in this paper and example reverse-correlation code can be found at [https://github.com/masseck]. Any additional information required to reanalyze the data reported in this paper is available from the lead contact upon request.

## EXPERIMENTAL MODEL AND SUBJECT DETAILS

### Animals

Adult male naïve wildtype (wt/wt), heterozygous (wt/-), and homozygous (-/-) NEX-Cre mice were bred on a C57BL/6J background using a NEX-Cre (wt/-) x NEX-Cre (wt/-) pairing ^1^. Male C57BL/6J mice (The Jackson Laboratory) were used as inter-strain control. All mice used in this study were between 2 and 9 months of age. Animals were maintained on a standard 12-hour light/dark cycle and housed in groups within individually ventilated cages (IVC, Zoonlab) under controlled conditions (22°C ± 2°C, 50% ± 5% humidity). Food and water were provided *ad libitum*. Mice were group-housed until the beginning of behavioral testing. All experiments were conducted during the dark phase, aligning with the animals’ primary activity period. Anxiety tests were performed with a one-week interval. All procedures followed the guidelines set by the Senator für Gesundheit, Frauen und Verbraucherschutz of the Freie Hansestadt Bremen.

## METHOD DETAILS

### Behavioral Experiments

#### Elevated Plus Maze (EPM)

The EPM is a widely used behavioral test to assess anxiety-like behavior, locomotor, and exploratory activity in rodents. This test leverages natural exploratory behavior of mice and their aversion to open and elevated areas. The EPM consists of two open arms (33.5 cm in length, 5 cm wide) and two closed arms (33.5 cm in length, 17 cm high wall, 5 cm wide) connected by a central platform, elevated 43 cm above the ground. Experiments were conducted under bright lighting conditions. Mice were individually transported from the animal facility to the experimental room and placed at the center of the maze, facing the open arm opposite to the experimenter. The behavior of each mouse was recorded for 5 min. Data were analyzed using EthoVision XT software (Noldus) with automated tracking quantifying the time spent in each zone, entrances to each zone, total distance and velocity.

#### Open Field Test (OFT)

The OFT is a commonly used behavioral assay to evaluate anxiety-like behavior, locomotion, and exploratory activity in rodents. The OFT was conducted under bright lighting conditions and takes place in a 50 cm (width) x 50 cm (depth) x 50 cm (height) Plexiglas arena. The arena was virtually divided into 16 equal squares, with the 4 inner squares representing the center zone (25 cm x 25 cm). Mice were individually transported from the mouse facility to the experimental room. Each mouse was placed in the lower left corner of the arena, facing the center. Behavior was recorded for 5 min and videos were analyzed using EthoVision XT software (Noldus). Automated tracking quantified the time spent in each zone, entrances to each zone, total distance and velocity. Time spent in the center zone and center zone entries are inversely correlated with anxiety levels. Circling behavior was tracked automatically using EthoVision XT software by counting body axis rotation which were defined as 360° Rotation around the axis of the center point and nose point. Grooming and rearing behaviors were analyzed offline. Rearing was defined as the mouse standing on its hind paws with forelimbs lifted off the ground.

#### Novelty Suppressed Feeding Test (NSFT)

For the NSFT, mice were transferred to a new cage and food-deprived for 24 hours prior to testing. The test was conducted in the open field arena, which was filled with approximately 200 g of fresh bedding per animal. A piece of filter paper (5 cm x 5 cm) with a standard food pellet was placed on the bedding in the center of the arena. Food pellets were weighed at the start of the experiment. The NSFT comprised three phases: habituation, test, and feeding. During the habituation phase, each animal was acclimated to the lighting conditions and the experimental room for 5 minutes. In the test phase, each mouse was placed in the back left corner of the open field arena, facing the food pellet. The test was terminated once the animal grasped the pellet with both front paws and began eating, with a maximum duration of 10 minutes allowed. The latency to begin eating was recorded. Following the test phase, each mouse was returned to its home cage and brought back to the mouse facility for the feeding phase, conducted immediately afterward. In this phase, each mouse had 5 minutes to eat the food pellet in the dark. The pellet was then weighed to calculate total food intake. After the experiment, all mice were given *ad libitum* access to food. The habituation and test phases were conducted under bright lighting conditions, while the feeding phase took place in darkness. Throughout the experiment, mice had *ad libitum* access to water. Latency to begin feeding is positively correlated with anxiety, while food intake serves as a control for normal feeding behavior.

#### Sucrose Preference Test (SPT)

The SPT is a two-bottle choice test used to assess anhedonic behavior, a core symptom of depression. This test is conducted in the home cage of each mouse during the active phase of the animals. For this experiment, 125 mL glass bottles were used, each equipped with a neck containing a small metal ball to prevent dripping during setup. The SPT consists of two phases: the habituation phase and the test phase. In the habituation phase, each mouse was given access to two bottles of tap water for 48 hours. Following this, the water bottles were replaced with two bottles containing freshly prepared 1% sucrose solution (w/v) for another 48 hours. During the test phase, each mouse had access to one bottle containing tap water and one bottle of 1% sucrose solution for 6 hours. Bottles were weighed before and after the test to calculate sucrose preference as a percentage of total liquid intake.

#### Social Interaction Test (SIT)

SIT is a behavioral assay used to assess social behavior and interactions in mice. This test was conducted in the open field arena (50 cm x 50 cm), where a small perforated Plexiglas chamber (10 cm wide x 6.5 cm deep x 42 cm high) was placed along one side of the arena. An area around this enclosure (12.5 cm x 25 cm) was defined as the social interaction area. The experiment consists of three phases: habituation, exploration, and interaction, following the protocol established by Golden and colleagues in 2011 ^38^. In the habituation phase, each mouse was individually brought to the experimental room and acclimated under red-light conditions for 1 hour. In the exploration phase, each mouse was placed at the center of the wall opposite the Plexiglas chamber in the open field arena and allowed to explore for 150 seconds before being returned to its home cage. For the interaction phase, an unfamiliar mouse (adult male CD-1 mouse) was placed in the Plexiglas chamber. Each mouse was placed again at the center of the wall opposite the chamber and allowed to explore the arena and interact with the unfamiliar mouse in the chamber for 150 seconds ^38^. The social interaction ratio (SI-ratio) is calculated by dividing the time spent in the interaction zone when the unfamiliar mouse is present by the time spent in the interaction zone when the unfamiliar mouse is absent. An SI-ratio of 1 indicates that the mouse spent equal time in the interaction zone during presence and absence of a social target. In addition, the time that each mouse spends in the interaction zone during the interaction phase was calculated as a percentage and given as the socialization time.

### Histology

#### Gelatine-Coated Microscope Slides

Microscope slides were polished, placed in a staining rack, soaked overnight in detergent and ddH₂O, rinsed, and air-dried at room temperature. The following day, the slides were cleaned in an ultrasonic bath with filtered isopropanol for 15 minutes, followed by immersion in boiling filtered 96% ethanol for 2 minutes. After drying at room temperature overnight, the slides were ready for gelatin coating. For coating, powdered gelatin was dissolved in ddH₂O to prepare a 0.5% solution (w/v) and heated to 70°C. Chromium potassium sulfate (CrK(SO₄)₂) was added to achieve a 0.05% solution (w/v), and the mixture was stirred until the color changed from yellow to green-blue. The solution was filtered and maintained at 75°C. For the first coating, cleaned slides were slowly dipped into the gelatin solution, covered and dried at room temperature overnight. The next day, a second coating was applied using freshly prepared gelatin-chromium potassium sulfate solution. The coated microscope slides were stored covered until use.

#### Perfusion and Golgi-Cox Staining

Following behavioral experiments, animals were euthanized via a lethal intraperitoneal injection of a ketamine/xylazine cocktail (130 mg/kg ketamine, 10 mg/kg xylazine). The mice were transcardially perfused with 1x phosphate-buffered saline (PBS) for 15 min, and their brains were stored for 24 hours at 4°C in a 30% sucrose solution (w/v) for cryoprotection.

Dendritic spine density was determined using Golgi-Cox staining, a widely used method for visualizing neuronal morphology, including dendritic spines. In this study the FD Rapid GolgiStain™ Kit (PK401, FD NeuroTechnologies, Ellicott City) was used, following the manufacturer’s instructions. After impregnation, the brains were snap-frozen at-70°C and stored at-80°C until the next day. On the following day, the brain tissue was mounted onto specimen discs by applying thin layers of distilled water using a paintbrush on dry ice. The brains were cut into 100 µm thick sections using a cryostat. Brain sections were mounted directly onto gelatin-coated glass slides and dried overnight at room temperature. The next day, the sections were stained according to the manufacturer’s instructions, coverslipped with Eukitt®, and stored at room temperature in the dark.

## QUANTIFICATION AND STATISTICAL ANALYSIS

All data analyses were performed in Python using the scikit-learn package ^40^.

### Behavioral Cluster Identification

#### PCA

PCA (Principal Components Analysis) is a linear statistical technique used for dimensionality reduction by transforming a dataset into a set of orthogonal principal components while preserving essential information and reducing noise and computational complexity. This transformation helps reveal underlying patterns, clusters, or relationships within the data, making PCA particularly valuable for preprocessing in machine learning tasks (e.g., SVM) to mitigate overfitting and enhance model performance. In this study, two principal components were derived from a dataset comprising behavioral parameters from the EPM and OFT (**Figure 1, Figure S 2+3**). The first principal component captures the direction of maximum variance in the data, representing the primary source of variability and illustrating the main axis of data distribution. The second principal component, orthogonal to the first, captures the next highest variance, reflecting the secondary axis of data spread. Together, these two components provide a simplified two-dimensional representation of the dataset while retaining most of its original variability, facilitating effective visualization. PCA was conducted using Python.

#### t-SNE Analysis

t-SNE (t-Distributed Stochastic Neighbor Embedding) is a non-linear dimensionality reduction technique designed for visualizing complex, high-dimensional datasets. In this study, the input dataset consisted of PCA-derived behavioral parameters from the EPM and OFT (Figure 1, Figure S 2+3). t-SNE operates by converting Euclidean distances between data points into conditional probabilities that indicate the likelihood of similarity between pairs of points. The algorithm then minimizes the divergence between the probability distributions of the high-dimensional data and its low-dimensional representation, ensuring that the resulting visualization accurately reflects local relationships within the data. This approach enables high-dimensional data to be represented in a low-dimensional space, making it easier to identify potential clusters or groups while preserving local structure ^39^. t-SNE was implemented using Python.

#### SVM

SVM (Support Vector Machine) is a supervised machine learning algorithm widely used for classification tasks. In this study, PCA was applied as a preprocessing step for dimensionality reduction, providing a compact feature set and serving as the basis for an optimized SVM with a Radial Basis Function (RBF) kernel. This approach enhances the model’s capacity to capture and process non-linear relationships within the data efficiently. SVM constructs optimal hyperplanes in the transformed feature space to distinguish between different behavioral classes. To ensure the generalizability and robustness of the model, 5-fold cross-validation was employed, partitioning the dataset into multiple folds for training and testing on various subsets. Additionally, a bootstrap method was used to generate multiple resampled datasets, enabling the estimation of the model’s variability and stability.

The combined use of PCA, SVM with RBF kernel, 5-fold cross-validation, and bootstrap resampling provides a rigorous framework for evaluating model performance, ensuring reliable and consistent behavioral classification outcomes. The aggregated confusion matrix summarizes classification results across all resampled datasets, offering insights into overall accuracy and the model’s effectiveness in distinguishing behavioral classes. The SVM analysis was performed using Python.

In the analysis using the SVM algorithm, a comprehensive set of behavioral parameters from the OFT and EPM were used as input features. From the OFT, locomotion metrics included total distance moved (cm) and mean velocity (cm/s), along with spatial preferences and anxiety-related behaviors such as frequency and total time spent in the border, corner, and center zones, as well as latency to first enter these zones. Additional metrics included frequency and total time of rearing, grooming, jumping, and body axis rotations. From the EPM, parameters included total distance moved (cm), mean velocity (cm/s), and exploratory measures such as closed and open arm entries, total time spent in closed and open arms, and latency to first enter these arms. The analysis also incorporated frequency and total time spent in the center field, along with latency to first enter the center. Together, these parameters provided a detailed behavioral profile, enabling the SVM algorithm to classify and interpret the data effectively (**Figure 1**, **Figure S 2+3**).

#### SHAP analysis

SHAP (SHapley Additive exPlanations) analysis was applied to interpret the feature contributions to the predictions made by the SVM model. This method offers detailed insights into feature importance and interactions, enhancing the interpretability of complex models, such as SVMs with RBF kernels. Using the SHAP library in Python, SHAP values were computed for each feature across all samples, enabling both a global assessment of feature impact and individualized explanations of specific predictions. This dual-level analysis provides a comprehensive understanding of how features influence the model’s outputs, making the decision-making process more transparent and interpretable.

## Data Analysis

All behavioral data were recorded using EthoVision XT software from Noldus. Data acquisition was performed automatically from recorded videos. Statistical analysis and graphical illustrations were created using GraphPad Prism 9 software. Microscope images of Golgi-Cox staining were processed for brightness and contrast using ImageJ. Quantification of dendritic spines was performed using ImageJ‘s dendritic spine counter plugin.

## Statistical Analysis

Normality was assessed with the Shapiro-Wilk test, and homogeneity of variances across groups was evaluated using Bartlett’s test. When both assumptions were met, a one-way ANOVA was performed to determine whether the means of independent groups differed, followed by Tukey’s post hoc test for pairwise comparisons. For non-parametric data, the Kruskal-Wallis ANOVA was used, followed by Dunn’s post hoc test for group comparisons. Outliers were identified using the ROUT method of regression, which combines a robust nonlinear regression with a False Discovery Rate-based test, employing a Q-value of 1% to estimate the likelihood of false positive results ^41^. Results are presented as violin plots with individual data points. Gray lines representing the 25th and 75th percentiles, solid black lines indicating the median. Significance levels were set at *p < 0.05, **p < 0.01, ***p < 0.001, and ****p < 0.0001.

