## Supplementary for "Uncovering Hidden Phenotypes in NEX-Cre Mice: Behavioral and Cellular Alterations Demand Re-Evaluation of a Widely Used Transgenic Line"

#### Supplementary Information

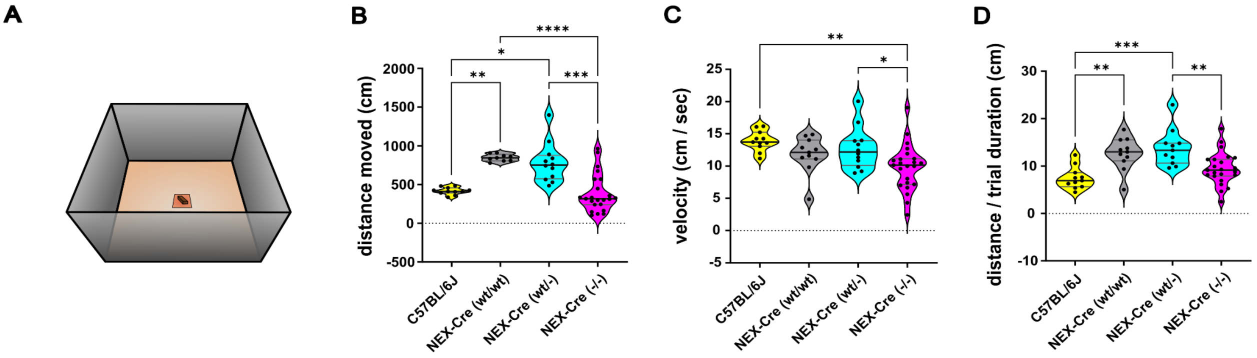


**Figure S 1 – Distance moved and Velocity in NSFT**. **(**A) Schematic overview of the NSFT. (B) NEX-Cre (-/-) covered significantly less distance compared to NEX-Cre (wt/wt) and NEX-Cre (wt/-) mice. NEX-Cre (wt/wt) and NEX-Cre (wt/-) covered significantly more distance compared to C57BL/6J (Kruskal Wallis ANOVA: H (3) = 28.98, p < 0.0001; Dunn`s post hoc: NEX-Cre (-/-) vs NEX-Cre (wt/wt) **** p < 0.0001, NEX-Cre (-/-) vs NEX-Cre (wt/-) ***p = 0.0005, NEX-Cre (wt/wt) vs C57BL/6J **p = 0.0067, NEX-Cre (wt/-) vs C57BL/6J *p = 0.0446). Sample sizes: C57BL/6J: n = 11, NEX-Cre (wt/wt): n = 9, NEX-Cre (wt/-): n = 12, NEX-Cre (-/-): n = 23. (C) NEX-Cre (-/-) showed significantly lower mean velocity than C57BL/6J and NEX-Cre (wt/-) (Ordinary one-way ANOVA: F (3,53) = 5.332, p = 0.0028; Tukey`s post hoc: C57BL/6J vs NEX-Cre (-/-) **p = 0.003, NEX-Cre (wt/-) vs NEX-Cre (-/-) *p = 0.046).Sample sizes: C57BL/6J: n = 11, NEX-Cre (wt/wt): n = 11, NEX-Cre (wt/-): n = 12, NEX-Cre (-/-): n = 23. (D) For normalization, the distance covered in the NSFT is divided by the respective latency to consume food. Relative to the trail duration, the NEX-Cre (-/-) have travelled less distance than the NEX-Cre (wt/-). The NEX-Cre (wt/wt) and the NEX-Cre (wt/-) have travelled more distance relative to the C57BL/6J (Ordinary one-way ANOVA: F (3,52) = 8.933, ****p < 0.0001; Tukey`s post hoc: C57BL/6J vs NEX-Cre (wt/wt) **p = 0.0034, C57BL/6J vs NEX-Cre (wt/-) ***p = 0.0002, NEX-Cre (wt/-) vs NEX-Cre (-/-) **p = 0.0043). Sample sizes: C57BL/6J: n = 11, NEX-Cre (wt/wt): n = 11, NEX-Cre (wt/-): n = 11, NEX-Cre (-/-): n = 23.

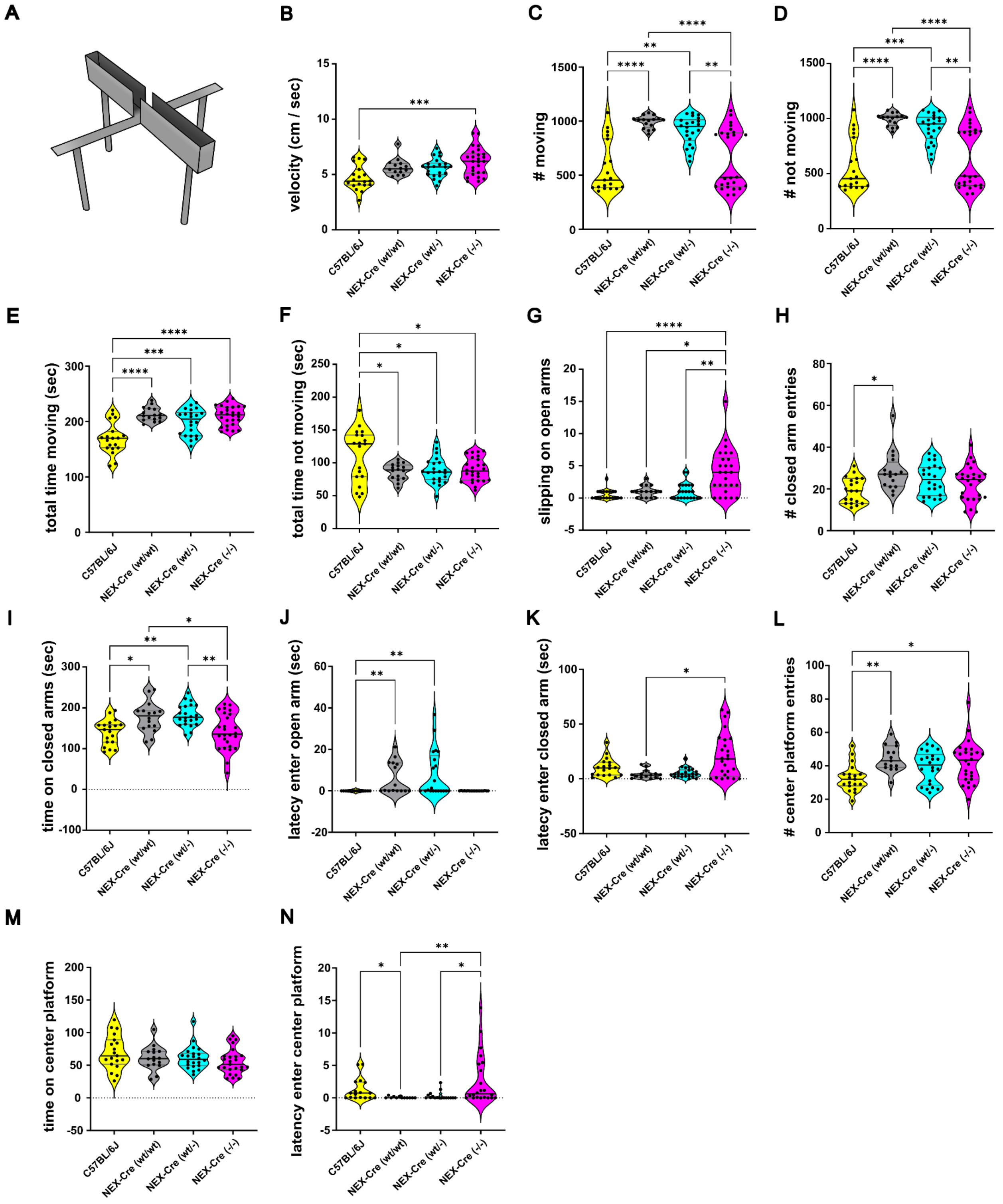


**Figure S 2 – Additional parameters measured in the EPM that were also included in the SVM analysis.** (A) Schematic overview of the EPM. (B) NEX-Cre (-/-) showed significantly increased mean velocity than C57BL/6J (Kruskal Wallis ANOVA: H (3) = 16.11, **p = 0.0011; Dunn`s post hoc: C57BL/6J vs NEX-Cre (-/-) *** p = 0.0004). Sample sizes: C57BL/6J: n = 19, NEX-Cre (wt/wt): n = 14, NEX-Cre (wt/-

): n = 22, NEX-Cre (-/-): n = 26. (C) NEX-Cre (-/-) showed a significantly lower movement frequency than NEX-Cre (wt/wt) and NEX-Cre (wt/-). NEX-Cre (wt/wt) and NEX-Cre (wt/-) showed a significantly higher movement frequency than C57BL/6J (Kruskal Wallis ANOVA: H (3) = 33.46, ****p < 0.0001; Dunn`s post hoc: NEX-Cre (-/-) vs NEX-Cre (wt/wt) **** p < 0.0001, NEX-Cre (-/-) vs NEX-Cre (wt/-) ** p = 0.0083, C57BL/6J vs NEX-Cre (wt/wt) **** p < 0.0001, C57BL/6J vs NEX-Cre (wt/-) ** p = 0.0011). Sample sizes: C57BL/6J: n = 19, NEX-Cre (wt/wt): n = 16, NEX-Cre (wt/-): n = 22, NEX-Cre (-/-): n = 26. (D) Consequently, NEX-Cre (-/-) showed a significantly lower frequency not moving than NEX-Cre (wt/wt) and NEX-Cre (wt/-). NEX-Cre (wt/wt) and NEX-Cre (wt/-) showed a significantly higher frequency not moving than C57BL/6J (Kruskal Wallis ANOVA: H (3) = 33.47, ****p < 0.0001; Dunn`s post hoc: NEX-Cre (-/-) vs NEX-Cre (wt/wt) **** p < 0.0001, NEX-Cre (-/-) vs NEX-Cre (wt/-) ** p = 0.0078, C57BL/6J vs NEX-Cre (wt/wt) **** p < 0.0001, C57BL/6J vs NEX-Cre (wt/-) *** p = 0.0010). Sample sizes: C57BL/6J: n = 19, NEX-Cre (wt/wt): n = 16, NEX-Cre (wt/-): n = 22, NEX-Cre (-/-): n = 26. (E) NEX-Cre (wt/wt), NEX-Cre (wt/-) and NEX-Cre (-/-) spent significantly more time moving compared to C57BL/6J mice. (Ordinary one-way ANOVA: F (3,79) = 17.29, ****p < 0.0001; Tukey`s post hoc: C57BL/6J vs NEX-Cre (wt/wt) ****p < 0.0001, C57BL/6J vs NEX-Cre (wt/-) *** p = 0.0002, C57BL/6J vs NEX-Cre (-/-) ****p < 0.0001). Sample sizes: C57BL/6J: n = 19, NEX-Cre (wt/wt): n = 16, NEX-Cre (wt/-): n = 22, NEX-Cre (-/-): n = 26. (F) Consequently, NEX-Cre (wt/wt), NEX-Cre (wt/-) and NEX-Cre (-/-) spent significantly less time not moving compared to C57BL/6J mice (Ordinary one-way ANOVA: F (3,79) = 4.420, **p = 0.0063; Tukey`s post hoc: C57BL/6J vs NEX-Cre (wt/wt) *p = 0.0196, C57BL/6J vs NEX-Cre (wt/-) * p = 0.0168, C57BL/6J vs NEX-Cre (-/-) *p = 0.0186). Sample sizes: C57BL/6J: n = 19, NEX-Cre (wt/wt): n = 16, NEX-Cre (wt/-): n = 22, NEX-Cre (-/-): n = 26. (G) NEX-Cre (-/-) were significantly more likely to slip their hind paws off the open arms compared to NEX-Cre (wt/wt), NEX-Cre (wt/-) and C57BL/6J mice (Kruskal Wallis ANOVA: H (3) = 23.29, ****p < 0.0001; Dunn`s post hoc: C57BL/6J vs NEX-Cre (-/-) **** p < 0.0001, NEX-Cre (wt/wt) vs NEX-Cre (-/-) * p = 0.0237, NEX-Cre (wt/-) vs NEX-Cre (-/-) ** p = 0.0034). Sample sizes: C57BL/6J: n = 19, NEX-Cre (wt/wt): n = 16, NEX-Cre (wt/-): n = 22, NEX-Cre (-/-): n = 26. (H) NEX-Cre (wt/wt) entered the closed arms significantly more often compared to C57BL/6J mice (Kruskal Wallis ANOVA: H (3) = 10.67, *p = 0.0136; Dunn`s post hoc: C57BL/6J vs NEX-Cre (wt/wt) * p = 0.0105). Sample sizes: C57BL/6J: n = 19, NEX-Cre (wt/wt): n = 16, NEX-Cre (wt/-): n = 22, NEX-Cre (-/-): n = 26. (I) NEX-Cre (-/-) spent significant less time on closed arms compared to NEX-Cre (wt/-) and NEX-Cre (wt/wt) while NEX-Cre (wt/-) and NEX-Cre (wt/wt) spent significantly more time on closed arms compared to C57BL/6J mice (Ordinary one-way ANOVA: F (3,79) = 7.976, ***p = 0.0001; Tukey`s post hoc: C57BL/6J vs NEX-Cre (wt/wt) *p = 0.0183, C57BL/6J vs NEX-Cre (wt/-) **p = 0.0025, NEX-Cre (-/-) vs NEX-Cre (wt/wt) *p = 0.0140, NEX-Cre (-/-) vs NEX-Cre (wt/-) **p = 0.0014). Sample sizes: C57BL/6J: n = 19, NEX-Cre (wt/wt): n = 16, NEX-Cre (wt/-): n = 22, NEX-Cre (-/-): n = 26. (J) NEX-Cre (wt/wt) and NEX-Cre (wt/-) had a significantly increased latency to enter the open arms compared to C57BL/6J mice. (Kruskal Wallis ANOVA: H (3) = 18.00, ***p = 0.0004; Dunn`s post hoc: C57BL/6J vs NEX-Cre (wt/wt) **p = 0.0058, C57BL/6J vs NEX-Cre (wt/-) **p = 0.0049). Sample sizes: C57BL/6J: n = 14, NEX-Cre (wt/wt): n = 15, NEX-Cre (wt/-): n = 21, NEX-Cre (-/-): n = 18. (K) NEX-Cre (-/-

) showed a significantly increased latency to enter closed arms compared to NEX-Cre (wt/wt) (Kruskal Wallis ANOVA: H (3) = 12.39, **p = 0.0062; Dunn`s post hoc: NEX-Cre (wt/wt) vs NEX-Cre (-/-) *p = 0.0124). Sample sizes: C57BL/6J: n = 18, NEX-Cre (wt/wt): n = 15, NEX-Cre (wt/-): n = 22, NEX-Cre (-/-): n = 25. (L) NEX-Cre (-/-) and NEX-Cre (wt/wt) showed significantly increased center platform entries compared to C57BL/6J mice. (Ordinary one-way ANOVA: F (3,78) = 4,737, **p = 0.0043; C57BL/6J vs NEX-Cre (wt/wt) **p = 0.0058, C57BL/6J vs NEX-Cre (-/-) *p = 0.0137). Sample sizes: C57BL/6J: n = 19, NEX-Cre (wt/wt): n = 15, NEX-Cre (wt/-): n = 22, NEX-Cre (-/-): n = 26. (M) All groups spent a similar amount of time on the center platform (Kruskal Wallis ANOVA: H (3) = 4,367, ns p = 0.2245): Sample sizes: C57BL/6J: n = 19, NEX-Cre (wt/wt): n = 16, NEX-Cre (wt/-): n = 22, NEX-Cre (-/-): n = 25. (N) NEX-Cre (-/-) had a significantly increased latency to enter the center platform compared to NEX-Cre (wt/wt) and NEX-Cre (wt/-) mice, while C57BL/6J had a significantly increased latency to enter the center platform compared to NEX-Cre (wt/wt) mice (Kruskal Wallis ANOVA: H (3) = 14.99, **p = 0.0018; Dunn`s post hoc: C57BL/6J vs NEX-Cre (wt/wt)

*p = 0.0498, NEX-Cre (wt/wt) vs NEX-Cre (-/-) **p = 0.0072, NEX-Cre (wt/-) vs NEX-Cre (-/-) *p = 0.0396).
Sample sizes: C57BL/6J: n = 17, NEX-Cre (wt/wt): n = 12, NEX-Cre (wt/-): n = 17, NEX-Cre (-/-): n = 22.

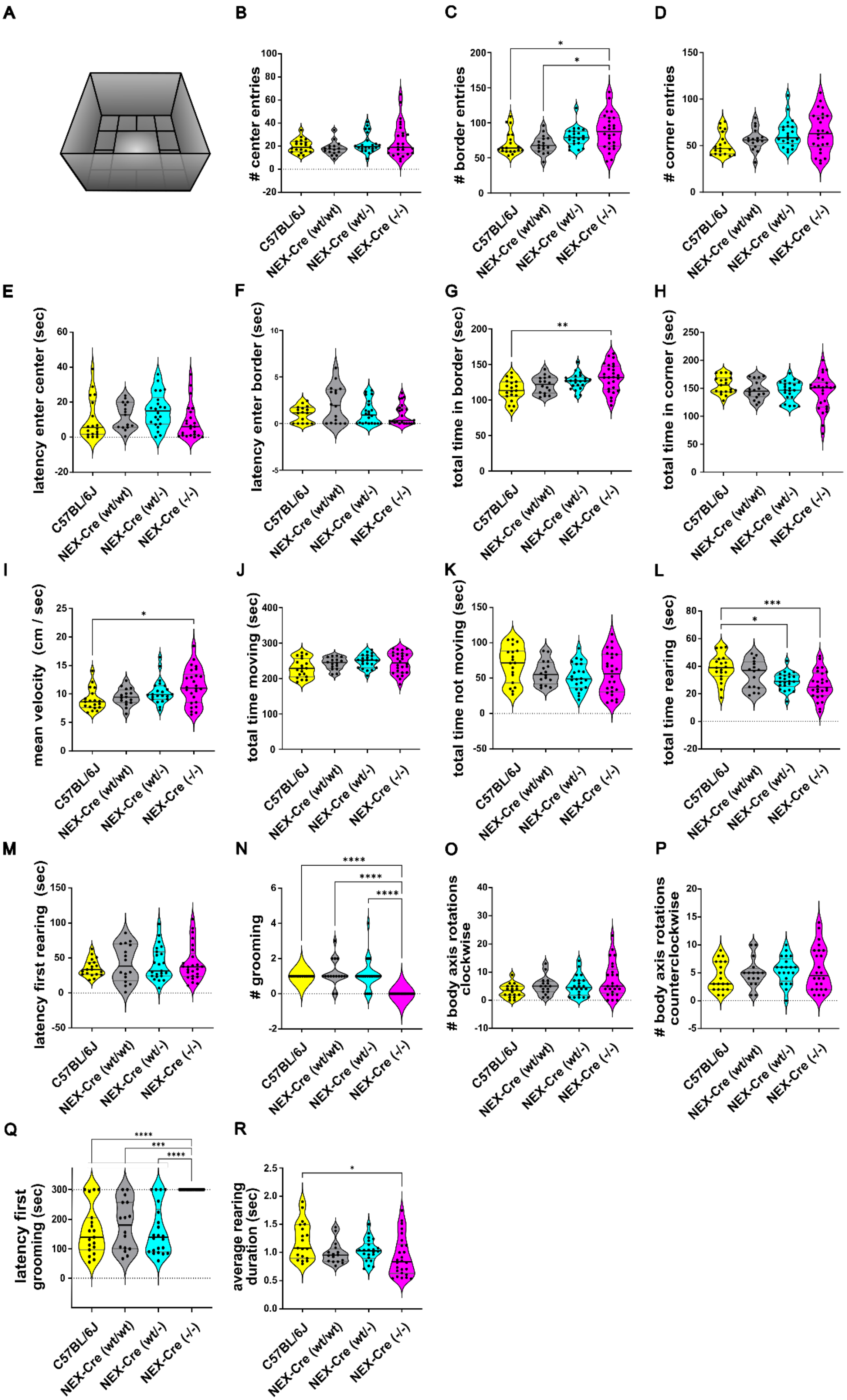
A
B
### center entries
C
### border entries
D
### corner entries
E
latency enter center (sec)
F
latency enter border (sec)
G
total time in border (sec)
H
total time in corner (sec)
I
mean velocity (cm / sec)
J
total time moving (sec)
K
total time not moving (sec)
L
total time rearing (sec)
M
latency first rearing (sec)
N
### grooming
O
### body axis rotations clockwise
P
### body axis rotations counterclockwise
Q
latency first grooming (sec)
R
average rearing duration (sec)
C57BL/6J
NEX-Cre (wt/wt)
NEX-Cre (wt/-)
NEX-Cre (-/-)

**Figure S 3 – Additional parameters measured in the OFT that were also included in the SVM analysis.** (A) Schematic overview of the EPM. (B) All groups entered the center field with similar frequency (Kruskal Wallis ANOVA: H (3) = 3.586, ns p = 0.3098). Sample sizes: C57BL/6J: n = 19, NEX-Cre (wt/wt): n = 16, NEX-Cre (wt/-): n = 22, NEX-Cre (-/-): n = 26.(C) NEX-Cre (-/-) entered significantly more often the border zone compared to NEX-Cre (wt/wt) and C57BL/6J mice (Kruskal Wallis ANOVA: H (3) = 12,47, ** p = 0.0059; Dunn`s post hoc: NEX-Cre (-/-) vs C57BL/6J *p = 0.02, NEX-Cre (-/-) vs NEX-Cre (wt/wt) *p = 0.0435). Sample sizes: C57BL/6J: n = 19, NEX-Cre (wt/wt): n = 16, NEX-Cre (wt/-): n = 21, NEX-Cre (-/-): n = 26. (D) All groups entered the corner zones with similar frequency (Kruskal Wallis ANOVA: H (3) = 7.511, ns p = 0.0573). Sample sizes: C57BL/6J: n = 19, NEX-Cre (wt/wt): n = 16, NEX-Cre (wt/-): n = 22, NEX-Cre (-/-): n = 26. (E) All groups entered the center field after a similar latency (Kruskal Wallis ANOVA: H (3) = 7.393, ns p = 0.0604). Sample sizes: C57BL/6J: n = 18, NEX-Cre (wt/wt): n = 15, NEX-Cre (wt/-): n = 22, NEX-Cre (-/-): n = 23. (F) All groups entered the border zone after a similar latency (Kruskal Wallis ANOVA: H (3) = 3.191, ns p = 0.3631). Sample sizes: C57BL/6J: n = 16, NEX-Cre (wt/wt): n = 16, NEX-Cre (wt/-): n = 22, NEX-Cre (-/-): n = 24. (G) NEX-Cre (-/-) spent significantly more time in border zones than C57BL/6J mice (Ordinary one-way ANOVA: F (3,79) = 5.323, **p = 0.0022; Tukey`s post hoc: NEX-Cre (-/-) vs C57BL/6J **p = 0.0012). Sample sizes: C57BL/6J: n = 19, NEX-Cre (wt/wt): n = 16, NEX-Cre (wt/-): n = 22, NEX-Cre (-/-): n = 26. (H) All groups spent similar time in corner zones (Ordinary one-way ANOVA: F (3,79) = 1.423, ns p = 0.2422). Sample sizes: C57BL/6J: n = 19, NEX-Cre (wt/wt): n = 16, NEX-Cre (wt/-): n = 22, NEX-Cre (-/-): n = 26. (I) NEX-Cre (-/-) had a significantly increased mean velocity compared to C57BL/6J mice (Kruskal Wallis ANOVA: H (3) = 10.33, * p = 0.0160; Dunn`s post hoc: NEX-Cre (-/-) vs C57BL/6J *p = 0.0157). Sample sizes: C57BL/6J: n = 19, NEX-Cre (wt/wt): n = 16, NEX-Cre (wt/-): n = 22, NEX-Cre (-/-): n = 26. (J) All groups spent similar time moving (Ordinary one-way ANOVA: F (3,79) = 2.12, ns p = 0.1043). Sample sizes: C57BL/6J: n = 19, NEX-Cre (wt/wt): n = 16, NEX-Cre (wt/-): n = 22, NEX-Cre (-/-): n = 26. (K) All groups spent similar time not moving (Ordinary one-way ANOVA: F (3,79) = 1.959, ns p = 0.127). Sample sizes: C57BL/6J: n = 19, NEX-Cre (wt/wt): n = 16, NEX-Cre (wt/-): n = 22, NEX-Cre (-/-): n = 26. (L) NEX-Cre (-/-) and NEX-Cre (wt/-) mice spent significantly more time rearing compared to C57BL/6J (Ordinary one-way ANOVA: F (3,79) = 7.212, ***p = 0.0002; Tukey`s post hoc: NEX-Cre (-/-) vs C57BL/6J ***p = 0.0002, NEX-Cre (wt/-) vs C57BL/6J *p = 0.0101). Sample sizes: C57BL/6J: n = 19, NEX-Cre (wt/wt): n = 16, NEX-Cre (wt/-): n = 22, NEX-Cre (-/-): n = 26. (M) All groups had a similar latency to start rearing (Kruskal Wallis ANOVA: H (3) = 0.2406, ns p = 0.9708). Sample sizes: C57BL/6J: n = 19, NEX-Cre (wt/wt): n = 16, NEX-Cre (wt/-): n = 23, NEX-Cre (-/-): n = 25. (N) NEX-Cre (-/-) exhibited a significantly reduced frequency of self-grooming events compared to C57BL/6J, NEX-Cre (wt/wt) and NEX-Cre (wt/-) mice (Kruskal Wallis ANOVA: H (3) = 41.82, ****p < 0.0001; Dunn`s post hoc: NEX-Cre (-/-) vs C57BL/6J ****p < 0.0001, NEX-Cre (-/-) vs NEX-Cre (wt/wt) ****p < 0.0001, NEX-Cre (-/-) vs NEX-Cre (wt/-) ****p < 0.0001). Sample sizes: C57BL/6J: n = 14, NEX-Cre (wt/wt): n = 16, NEX-Cre (wt/-): n = 22, NEX-Cre (-/-): n = 20. (O) All groups showed a similar frequency of body axis rotations in a clockwise direction (Kruskal Wallis ANOVA: H (3) =5.051, ns p = 0.1681). Sample sizes: C57BL/6J: n = 18, NEX-Cre (wt/wt): n = 16, NEX-Cre (wt/-): n = 22, NEX-Cre (-/-): n = 25. (P) All groups showed a similar frequency of body axis rotations in a counterclockwise direction (Kruskal Wallis ANOVA: H (3) = 2.165, ns p = 0.5388). Sample sizes: C57BL/6J: n = 19, NEX-Cre (wt/wt): n = 16, NEX-Cre (wt/-): n = 21, NEX-Cre (-/-): n = 24. (Q) Illustration of the latency in seconds until animals initiated self-grooming behavior during OFT. The total test duration was 300 seconds, and animals that did not exhibit grooming behavior are represented by data points at the 300-second mark. As no self-grooming was observed in NEX-Cre (-/-), the latency is significantly longer than in the other groups. There is no significant difference in the latency of first grooming between the other groups (Kruskal Wallis ANOVA: H (3) = 33.03, ****p < 0.0001; Dunn`s post hoc: NEX-Cre (-/-) vs C57BL/6J ****p < 0.0001, NEX-Cre (-/-) vs NEX-Cre (wt/wt) ***p = 0.0002, NEX-Cre (-/-) vs NEX-Cre (wt/-) ****p < 0.0001). Sample sizes: C57BL/6J: n = 19, NEX-Cre (wt/wt): n = 16, NEX-Cre (wt/-): n = 22, NEX-Cre (-/-): n = 20. (R) Compared to C57BL/6J mice, the NEX-Cre (-/-) spent significantly less time per rearing (Kruskal Wallis ANOVA: H (3) = 9.214, ****p = 0.0266; Dunn`s post hoc: NEX-Cre (-/-) vs C57BL/6J *p = 0.0201). Sample sizes: C57BL/6J: n = 19, NEX-Cre (wt/wt): n = 16, NEX-Cre (wt/-): n = 22, NEX-Cre (-/-): n = 20.
